# *Kelch13* and *MDR1* Polymorphisms, and Drug Effectiveness at Day 3 after Dihydroartemisinin-Piperaquine Treatment for *Plasmodium falciparum* Malaria on Bioko Island, Equatorial Guinea: 2014-2017

**DOI:** 10.1101/594366

**Authors:** Yu-Zhong Zheng, Jiang-Tao Chen, Xue-Yan Liang, Carlos Salas Ehapo, Urbano Monsuy Eyi, Hui-Ying Huang, Wei-Zhong Chen, Li-Yun Lin, Dong-De Xie, Yu-Ling Wang, Guo-Wei Chen, Xiang-Zhi Liu, Guang-Cai Zha, Huan-Tong Mo, Xin-Yao Chen, Jian Li, Ting-Ting Jiang, Min Lin

## Abstract

Artemisinin (ART) combination therapies were introduced on malaria endemic Bioko Island in 2004 through Bioko Island Malaria Control Project. Recently, ART-resistant *Plasmodium falciparum* strain with *Kelch13* (K13) propeller M579I mutation originating from Equatorial Guinea was observed as an increased parasite clearance time on day 3 after dihydroartemisinin-Piperaquine (DHA-PIP) treatment (D3 positivity). Here, we surveyed DHA-PIP effectiveness and molecular markers of drug resistance at D3 after DHA-PIP treatment on Bioko Island from 2014 to 2017. Among the 371 uncomplicated *P. falciparum* patients, 86.3% (320/471) were successfully followed up at D3. 5.9% (19/320) of patients showed D3 positivity. K13 and MDR1 gene were successfully sequenced from 46 patients collected at D0 (baseline population) and 19 D3-positivity patients. Five non-synonymous K13 mutations (H136N; K189N; K248N; K326E; K332N) were found. There was no statistical difference in the frequency of these K13 mutations between baseline population and D3-positivity samples (*p*>0.05). Additionally, none of the K13 propeller polymorphisms known to be involved in ART-resistance in Asia or Africa were detected. For MDR1 gene, 38.5% (25/65) carried N86Y mutation; 73.8% (48/65) the Y184F mutation. Parasites surviving DHA-PIP at D3 post-treatment were significantly more likely than the baseline population to carry the N86Y (*p* <0.05). These results suggest that K13 is not the best predictive molecular marker for ART resistance in Africa. More isolates from cases with delayed parasite clearance after DHA-PIP treatment indicated that *in vitro* and *in vivo* monitoring for ART derivatives and ACT partner drugs should be regularly performed on Bioko Island, Equatorial Guinea.

## INTRODUCTION

Malaria is one of the most important tropical parasitic diseases with an estimated 219 million cases and 435,000 deaths in 2017 (1). The emergence of *Plasmodium falciparum* resistance to anti-malarial drugs has been threatening the world’s malaria control and elimination efforts (2,3). Currently, World Health Organization (WHO) recommends artemisinin (ART) combination therapies (ACTs) as first-line treatment for uncomplicated *P. falciparum* malaria many endemic countries worldwide, and shown to have greatly contributed to the reduction in malaria morbidity and mortality (4). In Recent years, strong evidences supported that malaria parasites resistant to ARTs have emerged and are spreading in certain parts of Southeast Asia, and raises concerns that resistance may emerge and become widespread in high-burden settings, such as Africa (5,6). Untill now, the efficacy of ACTs remains high in Africa. However, evidence based on classical microscopic parasite detection suggests that a proportion of ACT-treated children in Kenya do not completely clear *P. falciparum* parasitemia (7). Additionally, recently Lu et al. observed (2017) an ART-resistant *P. falciparum* strain originating from Equatorial Guinea, Africa (6).

Characterization of ART-resistant parasite is primarily discriminated by nucleotide polymorphisms in the propeller domain of the *P. falciparum* kelch13 gene (K13, *Pf3D7_1343700*) (8). In Southeast Asia, K13 propeller polymorphism, such as Y493H, R539T, I543T and C580Y, is considered a reliable molecular marker for ART-resistant parasites, which showed increased survival rates in ring-stage survival assays (RSAs) and delayed parasite clearance in ACT-treated patients (5,8,9). Parasite clearance can be quantified either by the detection of parasites in patients on day 3 (i.e., “D3 positivity”) (10). The proportion of patients who are still parasitaemic on D3 after treatment with artesunate or ACT is currently an indicator for routine monitoring to identify suspected ART resistance (10).

Bioko Island, Equatorial Guinea, with historically high malaria transmission, has been subject to extensive interventions including intensive vector control, improved case management, intermittent preventative treatments (IPT) and behavioural change interventions since 2004 through the Bioko Island Malaria Control Project (BIMCP) (11). Through BIMCP, including the introduction of indoor residual spraying (IRS) and distribution of insecticide-treated nets (ITNs) to all households on Bioko Island, first-line artemisinin containing antimalarials (ACAs) free of charge, the parasite prevalence reduced from 43.3 to 10.5% between 2004 and 2016 (11). ACAs used in Equatorial Guinea have included artesunate plus sulfadoxine-pyrimethamine, artesunate plus amodiaquine and dihydroartemisinin-piperaquine (DHA-PIP) (12). Although ACAs are available and free, many people still sought their medicines at a private pharmacy rather than received treatment from the public health services (12). Additionally, falsified ACAs have been reported on Bioko Island, with the prevalence ranging between 6.1% and 16.1%, depending on the sampling method used (12). All of this increased the risk of ART-resistance in the region.

Our previous study showed that the absence of mutations in the propeller region of K13 in parasites from Bioko Island during 2013-2014 (13). The objectives of this study were to investigate the *K13* and *MDR1* polymorphisms, and drug effectiveness at Day 3 after DHA-PIP Treatment for *P. falciparum* Malaria on Bioko Island from 2014 to 2017.

## RESULTS

### General characters

From January 2014 to December 2017, we enrolled 471 patients with *P. falciparum* uncomplicated malaria from Bioko, Equatorial Guinea. The median (interquartile range [IQR]) age of patients was 26 years (18 to 38.5), and 36.9% of them were women. The median (IQR) parasite density was 3300 parasites/μl (1840 to 7000). Among the 471 enrolled patients, 86.3% (320/471) were successfully followed up on D3. The median (IQR) age of these patients was 26 years (17 to 37), and 40.0% of them were women. The median (IQR) parasite density was 3400 parasites/μl (1810 to 7940).

Among the 320 patients evaluated on D3, 5.9% (19/320) showed D3 positivity by microscopy, which was also confirmed and identified the *Plasmodium* species by PCR-HRM (14,15). These 19 patients had D0 and D3 parasite densities of 900 to 8000 and 40 to 2120 parasites/μl, respectively.

### Analysis of *K13* propeller gene

K13 propeller gene was successfully amplified and sequenced from 65 patients including baseline population (46 patients randomly selected from the 301 patients without ART-resistance, collected at D0) and 19 D3-positivity patients. Five non-synonymous mutations (H136N, CAT>AAT; K189N, AAG>AAC; K248N, AAG>AAT; K326E, AAA>GAA and K332N, AAA>AAC) were detected from the two groups (Table 1). However, none of the polymorphisms known to be involved in ART-resistance in Asia (8) or Africa (M579I) (6) were detected. There was no statistical difference in the frequency of these non-synonymous mutations between baseline population and D3 samples (*p*>0.05).

**Table 1.** Kelch13 and MDR1 polymorphisms of Bioko *P. falciparum* isolates in baseline population and D3-positively with DHA-PIP treatment

| Polymorphisms | Baseline population, % (n=46) | D3, % (n=19) | Odds ratio (95 % CI) | p-value |
| --- | --- | --- | --- | --- |
| ***K13* gene** |  |  |  |  |
| H136N(CAT>AAT) | 2 (4.3) | 0 (0.0) | 0.96 (0.90-1.02) | 0.356 |
| K189N(AAG>AAC) | 18 (39.1) | 7 (36.8) | 1.06 (0.35-3.20) | 0.863 |
| K248N(AAG>AAT) | 1 (2.2) | 0 (0.0) | 0.98 (0.94-1.02) | 0.517 |
| K326E(AAA>GAA) | 1 (2.2) | 0 (0.0) | 0.98 (0.94-1.02) | 0.517 |
| K332N(AAA>AAC) | 1 (2.2) | 0 (0.0) | 0.98 (0.94-1.02) | 0.517 |
| ***MDR1* gene** |  |  |  |  |
| N86Y(AAC>TAT) | 14 (30.4) | 11 (57.9) | 3.14 (1.04-9.50) | 0.038* |
| Y184F(TAT>TTT) | 31 (67.4) | 17 (89.5) | 4.11 (0.84-21.86) | 0.065 |

\*Baseline population: patients randomly selected from the 301 patients without ART-resistance, collected at D0; D3: the D3-positively patients after DHA-PIP treatment.

The *K* value for the whole gene of the 65 samples was 0.763. The highest nucleotide differences was found in the *Plasmodium*-specific region (codons 1-440) (*K*=0.732) while the lowest was found in the propeller domain (codons 441-726) (*K*=0.030). The overall haplotype diversity (Hd) and nucleotide diversity (π) in this gene were estimated to be 0.631±0.037 and 0.00039, respectively. Tajima’s D test, Fu and Li’s tests were also performed to analyze the natural selection in the Bioko *P. falciparum* K13 gene. The evidence of selection occurring on the K13 gene was not very conclusive as Tajima’s D was negative (−1.590, *p*>0.05), but the Fu & Li’s D&F was positive (D: −3.546, *p*<0.05; F −3.415, *p*<0.05) for the whole gene.

### Analysis of *MDR1* gene

All samples was successfully amplified and sequenced for MDR1 codons 86, 184, 1034, 1042, and 1246. Of the 65 samples, 38.5% (25/65) carried N86Y (AAC>TAT) mutation; 73.8% (48/65) the Y184F (TAT>TTT) mutation. No mutation was found at 1034, 1042 and 1246. As shown in Table 1, the proportion of patients carrying the mutation N86Y and Y184F in the baseline population was higher than that on D3. However, there was only a statistical difference in the proportion of N86Y mutation between the baseline population and D3-positive samples (*p* <0.05).

## DISCUSSION

There is a perennially high rate of malaria transmission throughout Equatorial Guinea, and DHA-PIP is currently commonly used for treatment. To the best of our knowledge, this is the first report to evaluate the effectiveness of routine DHA-PIP treatment for uncomplicated malaria, and described the drug-resistance genetic polymorphisms in D3-positivity *P. falciparum* isolates after DHA-PIP treatment in Equatorial Guinea. Our results showed that DHA-PIP drugs were efficacious in treating uncomplicated *P. falciparum* malaria in the region. This high cure rate of DHA-PIP for Bioko *P. falciparum* supports earlier findings from and across Africa (16-21). This ACT medicine is being increasingly recommended in malaria endemic countries as second-line treatment for *falciparum* malaria and also used for mass drug administration in special situations (21). However, it was found 5.9% (19/320) patients with D3 positivity, suggesting the possibility that a subset of parasites is becoming less susceptible to ART through mutation in some gene, that some individuals have relatively poor parasite-clearing immune responses, or that some patients were not self-administering their remaining doses of DHA-PIP. Based on these results, intensive surveillance of ART derivatives and ACT partner drugs must be conducted regularly on Bioko Island.

Due to the condition limitation, this study was not determining the *in vitro* susceptibility to ART derivatives using the ring survival test. Indeed, clinical resistance to ART is manifested by an increase in the ring-stage survival rate after contact with ART (5,8,9). In Southeast Asia, an increase in the ring-stage survival rate was also associated with the detection of K13 mutations including F446I, N458Y, Y493H, R539T, I543T, R561H and C580Y (5,8,9). In the present study, lower nucleotide differences (*K*=0.030) in the propeller domain of K13 gene were observed in the Bioko *P. falciparum* parasites collected during 2014 to 2017. No patients carried the mutations associated with ART-resistance in Asia (8). Notably, the K13 propeller mutation (M579I) associated with ART-resistance originating from Equatorial Guinea (6) was not observed on Bioko Island. These results are consistent with our previous study on the island in 2012-2014 (13), which revealed a limited number of genetic polymorphisms in the K13-propeller region. In addition, Bioko had the higher prevalence of isolates with mutant codons in the *Plasmodium*-specific region (46.2%, 30/65). The most frequently mutation was one non-synonymous mutation at proximal end (upstream region) K189N (83.3%, 25/30). Although the K189N mutation was observed at a comparatively higher frequency in the parasitic isolates, no correlation with clinical phenotype was observed among them (OR: 1.06, 95% CI 0.35-3.20, p=0.863) (Table 1). Previous study (22,23) revealed that the infections carried K189T/N had a median PC_1/2_ of 2.1 h (range 0.8-7.1) similar to that of infections with wild type parasites; 2.2 h (range 0.7-6.3). In Africa, significantly prolonged clearance has not yet been observed and the presently restricted variation in parasite clearance cannot be explained by K13 polymorphisms (23). All of these suggest that SNPs at K13 is not the best predictive molecular marker for ART-resistance among African patients. More isolates from cases of clinical failure or with delayed parasite clearance after treatment with ART derivatives are necessary to identify new molecular markers.

In our study, the frequency of N86Y mutation was significant higher in the patients prior to treatment (D0) than that on D3-positive patients (*p*<0.05). According to previous reports (24,25), the parasites containing the MDR1 86Y allele showed significantly higher piperaquine IC50s compared with those containing the MDR1 N86 allele. Multivariate analysis also revealed that MDR1 86Y allele was an associated factor of reduced piperaquine sensitivity. The important role of mutations in the MDR1 gene on *in vitro* piperaquine sensitivity has been confirmed by a recent study using genetically modified *P. falciparum* lines (26). Since the implementation of fixed-dose ACT through BIMCP, DHA-PIP would be started on Bioko Island; parasites with reduced piperaquine sensitivity might be selected in such areas. Thus, *in vitro* and *in vivo* monitoring should be regularly performed in the region.

In conclusion, these results suggest that K13 is not the best predictive molecular marker for ART-resistance in Africa. Although DHA-PIP remains efficacious in the study region and those ART-resistance mutations are not found, ACT may pose positive selection on the parasites. In additionally, the rate of transmission and the diversity of vector species on Bioko Island may also increase selection pressure on parasite strains (14). More isolates from cases with delayed parasite clearance after DHA-PIP treatment indicated that *in vitro* and *in viv*o monitoring for ART derivatives and ACT partner drugs should be regularly performed on Bioko Island, Equatorial Guinea.

## MATERIALS AND METHODS

### Study area

The study was carried out in the Malabo Regional Hospital and the clinic of the Chinese medical aid team to the Republic of Equatorial Guinea on Bioko Island. Ethical approval was obtained from the Ethics Committee of Malabo Regional Hospital. Bioko is an island 32 km off the west coast of Africa, and the northernmost part of Equatorial Guinea. It’s population, of approximately 334, 463 (2015 census, of which approximately 90% live in Malabo, the capital city) are at risk of malaria year-round (14). Before the launch of the BIMCP in 2004, entomological inoculation rates (EIR) on the island were in excess of 750 infectious bites per person per year (14).

### *Plasmodium falciparum* isolates

A total of 471 patients with uncomplicated malaria were collected and analysed between January 2014 and December 2017. Included patients were aged between 12 and 67 years, were residents on Bioko Island. Malaria patients were classified into uncomplicated malaria states according to the WHO criteria, which were defined as positive smear for *P. falciparum* and presence of fever (⩾37.5°C) (27). Consent was obtained from all participating subjects or their parents. Laboratory screening for malaria was done using an Immunochromatographic Diagnostic Test (ICT Malaria Combo Cassette Test) and confirmed using microscopic examination of blood smears (13,14). Extra blood drops were collected for a malaria smear and onto Whatman 903® filter paper (GE Healthcare, Pittsburgh, USA). The *Plasmodium* species were identified by a real-time PCR followed by high-resolution melting (PCR-HRM) as our previous reports (14,15). Patients positive for *P. falciparum* were treated with DHA-PIP according to the national treatment guidelines. The first dose was taken under the supervision of and observed by the pharmacy owner, who also provided the patients with clear explanation for consumption of the remaining two doses at home. All patients were asked again to confirm completion of DHA-PIP doses when they returned on D3 for clinical and parasitological follow-up.

### PCR amplification of K1 and MDR1 gene

Genomic DNA was extracted from dried filter bloodspots (DBS) with Genomic DNA Extraction Kit for Dry Blood Spot [(No. DP334) TIANGENE Biotech (Beijing) CO., Ltd], and following the manufacturer’s protocol.

We developed a new PCR method to amplify the entire K13 gene (Pf3D7_1343700) (Fig. 1A). The primary PCR primers (K13_PCR_Fand K13_PCR_R) amplified the expected 2,097 bp product under the following conditions: 95°C for 3 min; 30 cycles of 98°C for 10 s, 58°C for 10s, 72°C for 2 min; and final extension at 72°C for 10 min; The nested PCR primers (K13_N1_F and K13_N2_R) amplified the expected 2,027bp product under the following conditions: 95°C for 3 min; 35 cycles of 98°C for 10 s, 55°C for 5s, 72°C for 2 min; and final extension at 72°C for 10 min. TaKaRa Taq(tm) HS Perfect Mix (TaKaRAa, Carlsbad, CA) was used as the master mix and was supplemented with a 0.2 μM concentration of each primer. Final volumes for primary and nested PCRs were 25 μl (12.5 μl master mix plus 1 μl DNA template) and 50 μl (25 master mix plus l μl primary PCR product), respectively.

**FIG 1.**
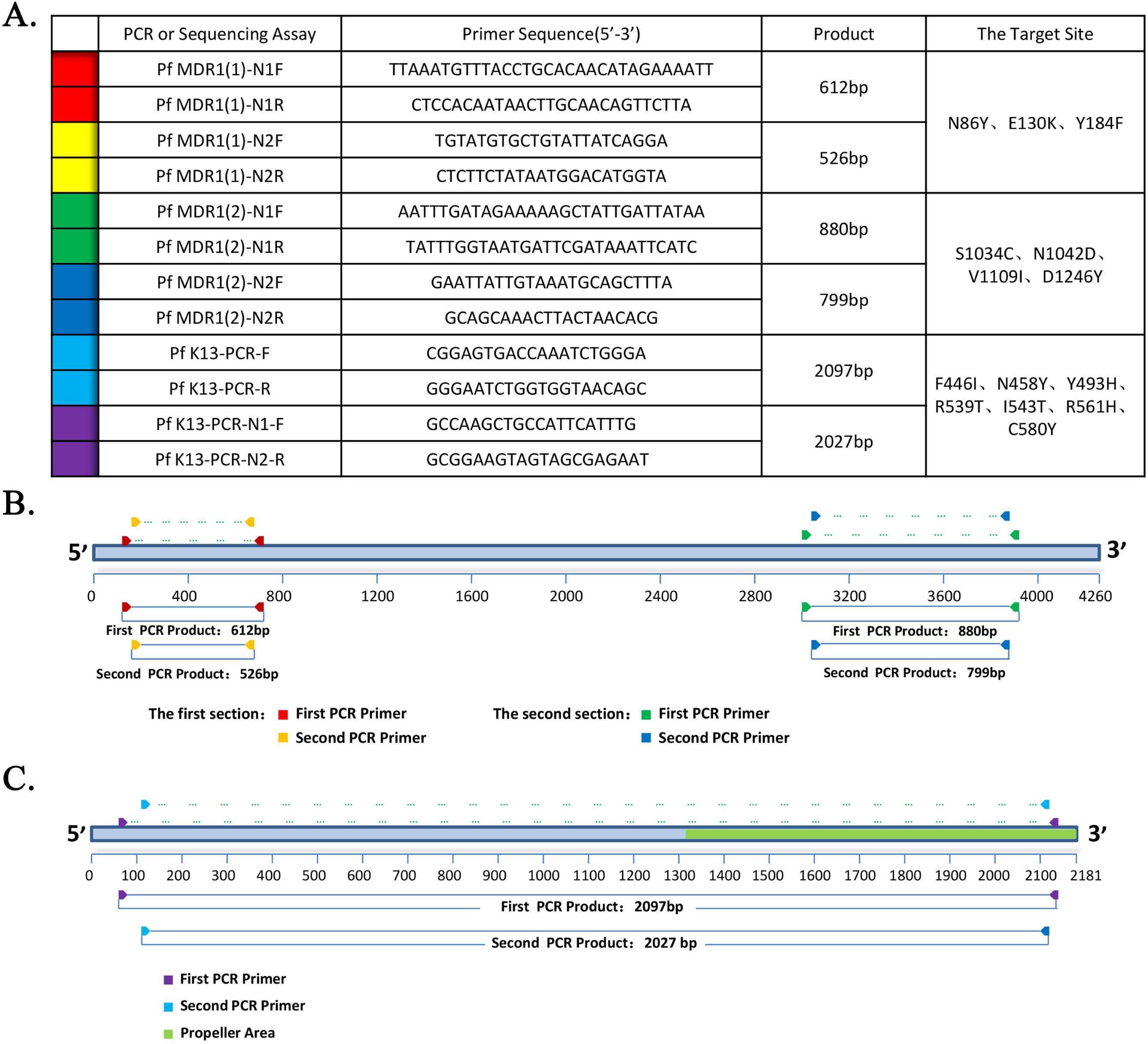
Primers and schematic representations of *K13* and *MDR1* nested PCR and sequencing strategies. New nested PCR and sequencing primers (A) were developed to capture the 7 MDR1 SNPs (B) and the whole K13 gene (C). The primers in the table are color-coded to match their positions in the schematics.

In order to capture MDR1 SNPs at codons 86, 130, 184, 1034, 1042, 1109, and 1246, a nest PCR were performed to amplify 2 shorter fragments (Fig. 1B) as our previous reports (28,29).

### Sequence polymorphism analysis

All PCR products were analyzed using 1.0% agar gel electrophoresis and DNA sequencing using an ABI 3730XL automated sequencer (PE Biosystems, CT, USA). The nucleotide and deduced amino acid sequences of K13 and MDR1 were analysed using EditSeq and SeqMan in the DNASTAR package (DNASTAR, Madison, WI, USA). The K13 and MDR1 sequences of the laboratory-adapted *P. falciparum* strain 3D7 (XM_001351086) was included in the alignment for comparison as a reference sequence. In K13 gene, the values of segregating sites (S), the average number of pair-wise nucleotide differences (*K*), haplotype diversity (Hd), and nucleotide diversity (π) were calculated using DnaSP version 5.10.00. The π was also calculated on a sliding window plot of 10 bases with a step size of 5 bp in order to estimate the stepwise diversity across the sequences. Tajima’s D test, Fu and Li’s D and F statistics analysis were performed using DnaSP package ver. 5.10.00 in order to evaluate the neutral theory of natural selection. The recombination parameter (R), which included the effective population size and probability of recombination between adjacent nucleotides per generation, and the minimum number of recombination events (Rm) were analysed using DnaSP version 5.10.00. The frequency data was analyzed using SPSS 17.0 (SPSS Inc., Chicago, IL). The *p*-value< 0.05 was considered statistically significant.

### Contributions

ML, JTC and YZZ designed the study. CSE, UME, DDX, YLW, GWC collected the samples, entered the data and validated microscopy. XYL, HYH and YZZ analyzed and interpreted the data. XZL, GCZ, HTM, XYC, JL, TTJ,WZC and LYL conducted the laboratory work (*P. falciparum* PCR and analysis of molecular markers). ML and YZZ wrote the paper. All authors critically reviewed the paper and approved the final version of the paper for submission.

### Declaration of Interest

The Authors report no conflicts of interest.

## Acknowledgments

The authors thank the Department of Health of Guangdong Province and Department of Aid to Foreign Countries of Ministry of Commerce of People’s Republic of China for their help. The authors also thank Santiago-m Monte-Nguba for his technical help during the samples collection and diagnosis. This work was partially supported by Natural Science Foundation of Guangdong Province (Grant No. 2016A03031311 to Jiang-Tao Chen; 2018A030307074 to Yu-Zhong Zheng) and Guangdong Science and Technology Project (Grant No. 2016A030303064 to Guang-Cai Zha).

